# Bitesize bundles F-actin and influences actin remodeling in syncytial *Drosophila* embryo development

**DOI:** 10.1101/2023.04.17.537198

**Authors:** Anna R. Yeh, Gregory J. Hoeprich, Anthony McDougal, Bruce L. Goode, Adam C. Martin

**Author notes:** Corresponding author: Adam C. Martin.

## Abstract

Actin networks undergo rearrangements that influence cell and tissue shape. Actin network assembly and organization is regulated in space and time by a host of actin binding proteins. The *Drosophila* Synaptotagmin-like protein, Bitesize (Btsz), is known to organize actin at epithelial cell apical junctions in a manner that depends on its interaction with the actin-binding protein, Moesin. Using RNAi, we showed that Btsz functions at earlier, syncytial stages of *Drosophila* embryo development. Btsz is required to stabilize pseudo-cleavage furrows that prevent metaphase spindle collisions and nuclear fallout prior to cellularization. While previous studies have focused on Btsz function through Moesin, we find that phosphorylated Moesin localized to the nuclear envelope and was not enriched at pseudo-cleavage furrows, suggesting a Moesin-independent function for Btsz in syncytial embryos. Consistent with this, mutants that affected all Moesin binding domain isoforms did not recapitulate pan-isoform Btsz depletion and we find that the C-terminal half of Btsz cooperatively binds to and bundles F-actin. We propose that Synaptotagmin-like proteins directly regulate actin organization during syncytial *Drosophila* development.

## Introduction

During animal development, actin networks are tightly regulated and undergo dynamic rearrangements to promote cell and tissue morphogenesis (Camuglia et al., 2021; Lecuit and Yap, 2015; Sokac et al., 2023). However, the range of mechanisms that control actin network assembly and organization during development is still not fully understood. Actin networks must be regulated at four major levels. First, is the control of assembly. The nucleation and assembly of monomeric G-actin (globular actin) into polymeric F-actin (filamentous actin) are tightly regulated in both time and space.

Spontaneous nucleation of actin is inhibited by many factors *in vivo* (Plastino and Blanchoin, 2019). Therefore, actin nucleators and polymerizers such as the formin, Diaphanous (Dia), as well as the Arp2/3 complex are essential in driving actin network formation. Dia assembles actin structures composed of linear (unbranched) filaments, whereas Arp2/3 complex assembles branched actin filament networks. Next, is the spatial organization of filaments into higher-order structures, or networks, which involves a plethora of actin binding and bundling/crosslinking proteins such as Fascin/Singed, Filamin/Cheerio and Fimbrin (Cant et al., 1994; Davidson et al., 2019; Krueger et al., 2019; Tilney et al., 2000). These networks are important for structure, transport, and transducing mechanical forces across the organism. Crosslinking actin filaments not only governs the network’s architecture but also its rigidity and contractile properties (Koenderink and Paluch, 2018). Third is the regulated disassembly or turnover of actin networks, which is orchestrated by a team of actin binding proteins that includes cofilin, AIP1, coronin, twinfilin, and cyclase-associated protein (CAP) (Goode et al., 2023). Finally, the cortical actin network must be connected to the cell plasma membrane through actin-binding proteins in order to provide structure and help induce cell shape changes (Chugh and Paluch, 2018). The ERM (Ezrin Radixin Moesin) family of proteins plays an essential role in this process (Algrain et al., 1993; Gould et al., 1989). When activated by phosphorylation through kinases, ERM proteins undergo a conformational change, exposing an F-actin binding and membrane protein-binding site through which it tethers F-actin to membranes (Gary and Bretscher, 1995; Zaman et al., 2021). Therefore, each cell must coordinate actin nucleation, assembly, and disassembly with higher-order actin network organization and membrane-to-cortex attachment.

The early *Drosophila* embryo provides an optimal system in which to study actin regulation due to its dynamic actin network rearrangements. At the earliest stages of development, the embryo is a syncytium consisting of one cell with multiple nuclei in a shared cytoplasm. The syncytium undergoes multiple rounds of synchronous modified cell divisions lacking cytokinesis and intervening gap phases, called nuclear divisions (Figure 1A) (Foe et al., 2000). Nuclear cycles (NC) 1-9 occur in the yolk at the center of the embryo, and nuclei migrate to the apical surface of the embryo for NC 10-13 (Foe et al., 2000). Centrosomes then help direct the formation of F-actin structures called ‘caps’ (Stevenson et al., 2002, 2001), which are positioned above the syncytial nuclei. During each nuclear cycle, F-actin assembly drives the growth of actin caps that expand into one another (Figure 1A). Collisions between neighboring caps lead to the formation of pseudo-cleavage furrows, structures made up of both actin and membrane that serve to compartmentalize mitotic spindles (Zhang et al., 2018) (Figure 1A). Pseudo-cleavage furrows act as a barrier between dividing spindles and are essential for proper nuclear divisions. Perturbing pseudo-cleavage furrows with drugs or depleting actin regulatory proteins often results in spindle collisions (Afshar et al., 2000; Callaini et al., 1992; Spracklen et al., 2019; Sullivan et al., 1993; Webb et al., 2009; Zallen et al., 2002).

**Figure 1.**
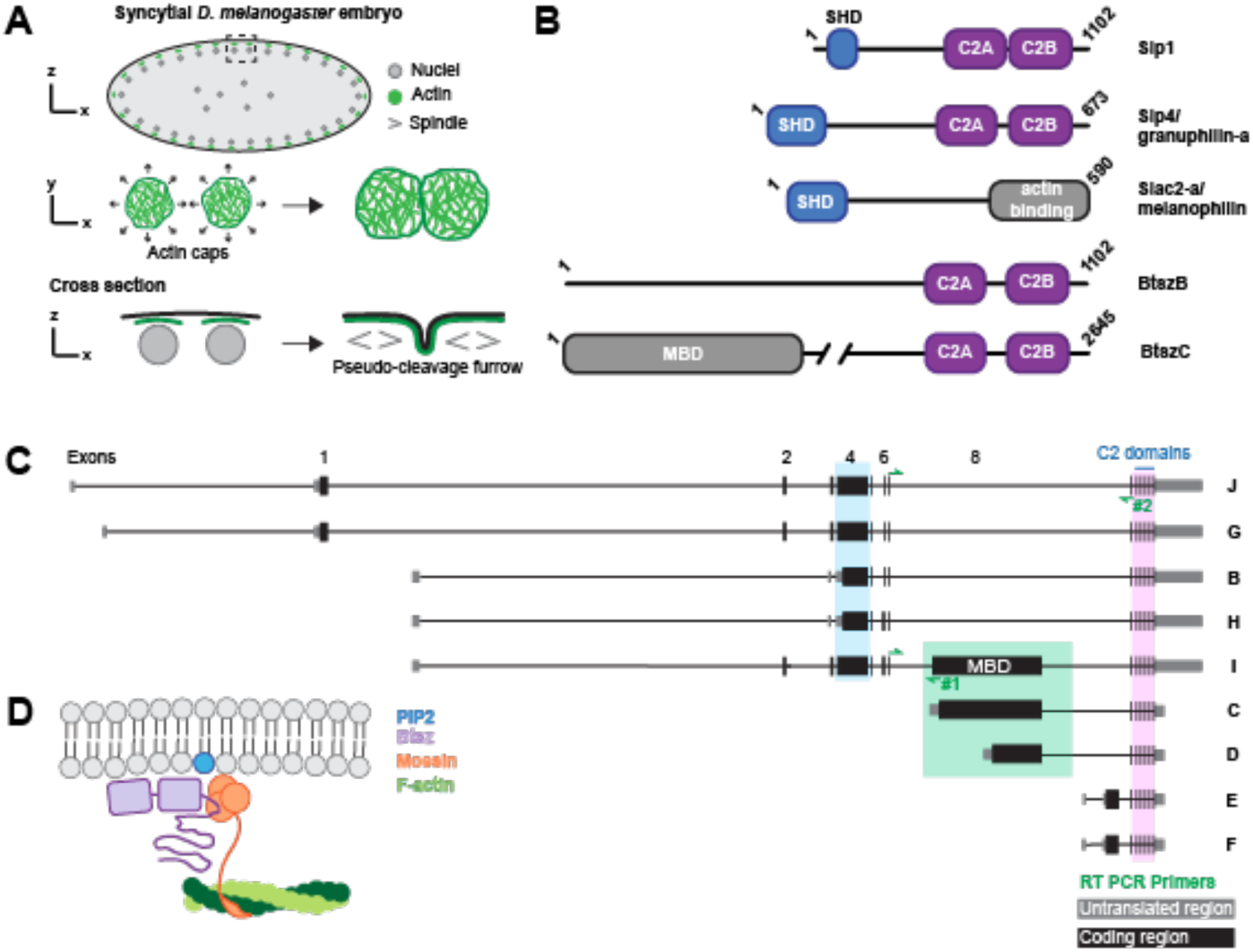
Btsz is a Synaptotagmin-like protein (Slp) with multiple splice isoforms. **(A)** Cartoon depiction of a syncytial embryo during a nuclear cycle. Sagittal plane through a syncytial embryo (top row). Zoomed in view of the actin caps in the dashed rectangular box where actin caps grow and collide with neighboring actin caps (middle row). Cross section of the zoomed in view where pseudo-cleavage furrows form from actin caps that have collided (bottom row). **(B)** Schematic of mammalian (mouse) Slp1, Slp4, Slac2-a, and fly Btsz isoform B and C proteins. **(C)** Schematic of the nine splice isoforms of the Btsz protein. Letters on the left-hand side denote the isoform name. Location of isoform-specific RT-PCR primer pairs #1 and #2 for non-MBD (BtszJ, G, B, and H) and BtszI, respectively, are shown with green arrows. **(D)** Model depicting Btsz’s interaction with the plasma membrane and Moesin. Btsz and Moesin localized to apical membrane through interactions with PIP2. There, Btsz can bind the FERM domain of Moesin and Moesin binds F-actin.

In addition to actin assembly, membrane trafficking is essential for syncytial blastoderm development. When membrane trafficking is perturbed by knocking down components of the trafficking machinery, such as Nuf and Rab11, pseudo-cleavage furrows are shorter and resemble those in embryos defective in actin assembly (Cao et al., 2008; Holly et al., 2015; Riggs et al., 2003; Rikhy et al., 2015; Rothwell et al., 1998; Sherlekar et al., 2020). Together, these findings indicate that actin network reorganization is coupled with membrane remodeling to create pseudo-cleavage furrows. However, there is still much to uncover about the roles of actin and/or membrane remodelers play in this process.

Synaptotagmin-like proteins (Slps) are potential candidates for coupling actin and membrane remodeling during syncytial blastoderm development. Slps are conserved across vertebrates and many invertebrates. Vertebrate Slps are characterized by an N-terminal Slp homology domain (SHD) and two C-terminal C2 domains (Figure 1B). The mammalian SHD interacts with the GTPase Rab27, and the C2 domain consists of the C2A and C2B domains that bind membrane, sometimes in a calcium-dependent manner (Fukuda, 2002; Gálvez-Santisteban et al., 2012). Specifically, the C2A domain binds with high affinity to PI(4,5)P_2_ or PI(3,4,5)P_3_ and the C2B domain binds with high affinity to PI(4,5)P_2_ (Gálvez-Santisteban et al., 2012; Lyakhova and Knight, 2014). A family of proteins related to Slps are Slac2s, Synaptotagmin-like proteins lacking C2 domains, which contain the SHD at their N-terminus but contain a unique C-terminus lacking the C2 domains (Figure 1B). Slac2 proteins function as adaptor proteins for cargo transport, interacting with Rab27A via their SHD, as well as Myosin Va via their myosin-binding sites found in the middle of Slac2 (Fukuda et al., 2002). Interestingly, the C-termini of Slac2-a and Slac2-c also bind G-and F-actin (Fukuda et al., 2002). Both Slps and Slac2s are Rab27 effectors important for vesicle secretion. For example, Slp4 is necessary for proper exocytosis and granule docking in the pancreas (Mizuno et al., 2016). During *de novo* epithelial formation, Slp2-a is required for targeting vesicles to the apical surface in a PIP_2_ dependent manner, and Slp4-a couples these vesicles to apical plasma membrane to generate lumen (Gálvez-Santisteban et al., 2012). Overall, there are six Slp genes and three Slac2 genes in mammals.

*Drosophila* encodes a single Slp gene, *Bitesize* (*Btsz*) that plays a role in actin cytoskeleton organization (Pilot et al., 2006b) similar to mammalian Slp4/granuphilin. Pilot et al. showed that Btsz is essential for maintaining adherens junctions and epithelial integrity. They discovered that Btsz binds phosphoinositides and that certain Btsz isoforms interact with the membrane-cortex crosslinker, Moesin (Moe), via a Moesin binding domain (MBD) (Figure 1C, D). During tracheal morphogenesis, Btsz recruits Moesin to the luminal membrane to promote proper actin organization (JayaNandanan et al., 2014). Therefore, previous work has established the importance of Btsz in regulating the actin cytoskeleton through Moesin. However, alternative splicing generates nine different Btsz isoforms and most of them lack the exon that encodes the MBD (Figure 1C). This raises the question of whether Btsz has additional, distinct functional roles in development.

In this study, we discovered a role for Btsz in regulating pseudo-cleavage furrow morphology and stability during syncytial stages of *Drosophila* embryo development, which in turn impacts spindle and nuclear morphology. This function of Btsz is mediated, at least in part, through functions beyond Moesin recruitment. We demonstrated this by examining phospho-Moesin localization, comparing genetic alleles that affect distinct sets of Btsz isoforms, and examining isoform-specific expression in the early embryo. We found that the C terminus of Btsz binds to F-actin and bundles F-actin *in vitro*, suggesting a direct role for Btsz in regulating actin architecture.

## Results

### Btsz is required for actin remodeling during syncytial blastoderm development

To examine Btsz function in early development, we used an RNAi line to maternally deplete embryos of Btsz (Figure S1A). This shRNA targets a sequence in the 3’ UTR shared by all isoforms, while primers used for RT-PCR detected MBD and Exon 4 containing isoforms (Figure S1A and B). Past work on Btsz focused on isoforms containing the Moesin-binding domain (MBD, Exon 8) and their roles during gastrulation in regulating actin organization, epithelial integrity, and lumen formation (JayaNandanan et al., 2014; Pilot et al., 2006b). Upregulation of Btsz isoforms containing the MBD at the mid-blastula transition was noted in previous studies (Pilot et al., 2006a). Knock-down of all Btsz isoforms revealed striking defects in syncytial embryo development as well as defects after epithelium formation, during gastrulation. We found that 60% of Btsz-RNAi embryos displayed syncytial or cellularization defects compared to 13% of control embryos and 92% of Arp3 RNAi embryos, which we used as a positive control (Figure S1C). Because defects prior to gastrulation in Btsz-depleted embryos have not been described, and the syncytial blastoderm is an excellent system in which to study actin remodeling, we further investigated Btsz function in the syncytial blastoderm.

To determine Btsz function in syncytial stages of embryonic development, we visualized actin in live embryos by driving mCherry::MoesinABD (mCh::Moe) expression with the maternal Gal4 driver to observe actin cap and pseudo-cleavage furrow structure and dynamics (Xie et al., 2021). We found that Btsz depletion affected the organization of F-actin compartments above the nuclei (Figure 2A; Video 1). We measured nuclear F-actin compartment size once the caps had finished expanding in control and Btsz-RNAi embryos. We quantified cycle 12 compartments because of good nuclear spacing and it has a high number of caps without the caps being too crowded.

**Figure 2.**
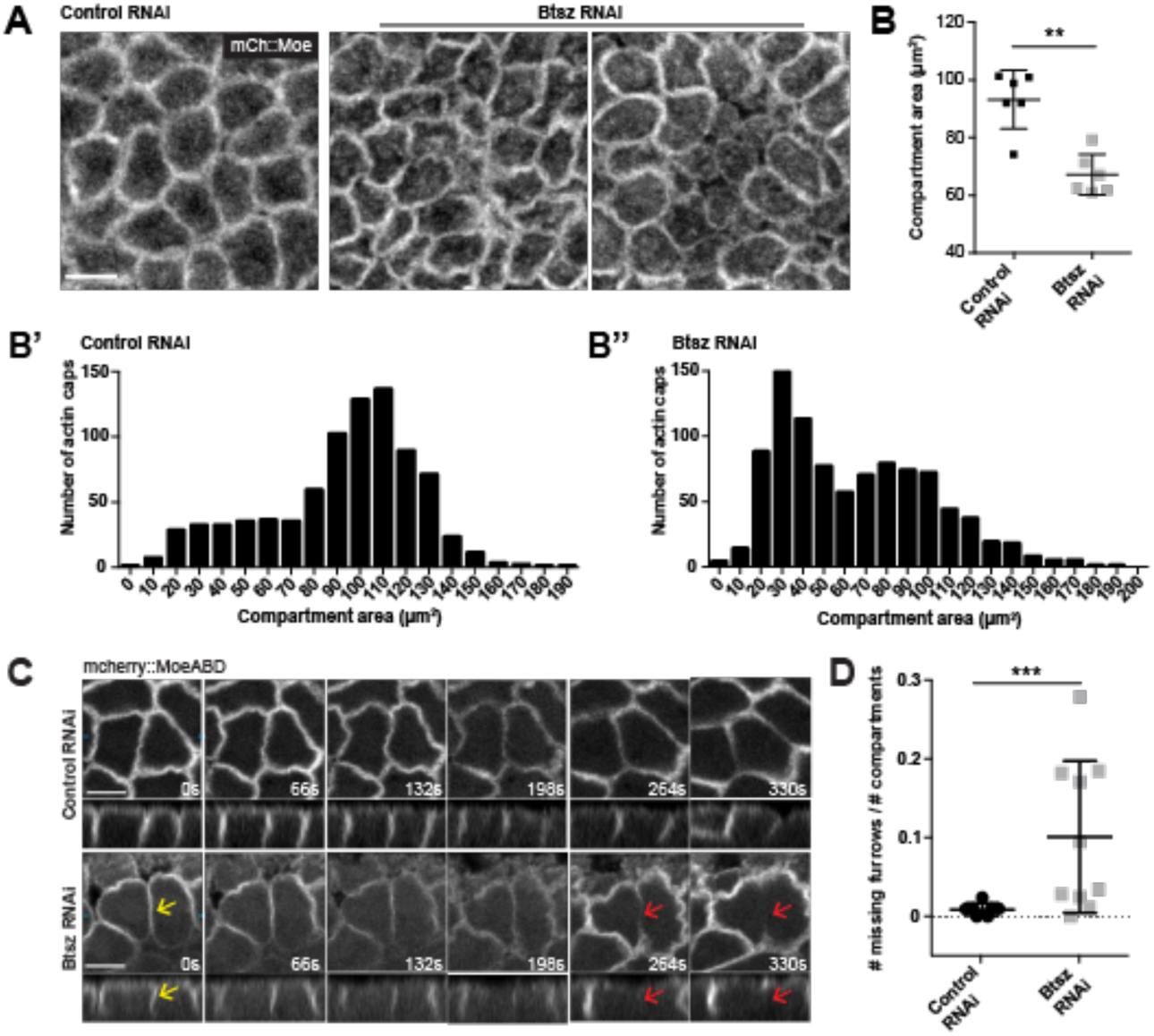
Btsz is required for actin remodeling during syncytial blastoderm development. **(A)** Maximum projection of a top-down view of control and Btsz RNAi nuclear cycle 12 actin caps. Actin was visualized live using the mCherry::MoesinABD marker. **(B)** In Btsz-depleted embryos, the average cap size is smaller compared to those in the control. Each data point is an average of all caps in the field of view per embryo. n = 6 embryos for both control RNAi and Btsz RNAi. p = 0.0043, Mann-Whitney U test. **(B’, B’’)** Histogram of raw data of actin cap area from Figure 2B. There is a peak of small actin caps (∼30 μm^2^) in Btsz RNAi embryos that is absent in the control embryos. **(C)** Time-lapse showing the formation of pseudo-cleavage furrows in control and Btsz-depleted embryos during nuclear cycle 12, with single slices (top panels) and orthogonal views (bottom panels). Actin was visualized live using the mCherry::MoesinABD marker. In Btsz RNAi embryo, a pseudo-cleavage furrow forms normally (yellow arrows), but breaks and recedes (red arrows) during metaphase while neighboring furrows are at their maximum length. **(D)** Number of missing furrows normalized over total number of actin caps during nuclear cycle 12. In Btsz-depleted embryos, pseudo-cleavage furrows prematurely recede during mitosis more often than in wild type embryos. Each point is the ratio of missing furrows to actin caps in a field of view (212 x 106 μm) per embryo. n = 7 embryos for control RNAi, n = 10 embryos for Btsz RNAi, p = 0.0007, Mann-Whitney U test. Scale bars are 10 µm.

Our results show that in Btsz-depleted embryos, the average compartment area is smaller compared to those of control embryos (Figure 2A-B). When comparing the distribution of compartment areas, there is a bimodal distribution of compartment sizes and a greater proportion of small compartments (<80 µm^2^) in Btsz-RNAi embryos compared to the control (Figure 2B-B’’).

Actin cap expansion and subsequent collisions between neighboring caps drive pseudo-cleavage furrow formation and growth (Zhang et al., 2018) so we hypothesized that pseudo-cleavage furrows would be affected in Btsz-RNAi embryos. Interestingly, small actin caps in Btsz-depleted embryos still formed pseudo-cleavage furrows between nuclei. Live imaging of actin in pseudo-cleavage furrows did reveal that a higher frequency of furrows prematurely retract during mitosis in Btsz-RNAi embryos (Figure 2C, D; Video 2). These unstable furrows formed normally (Figure 2C, yellow arrows) but receded during metaphase, resulting in furrows that are either very short or no longer visible (Figure 2C, red arrows). Because furrow regression could reflect a membrane trafficking defect (Holly et al., 2015; Riggs et al., 2003; Rikhy et al., 2015; Rothwell et al., 1998), we tested whether Btsz-RNAi affected ER or golgi structure.

Btsz-RNAi did not affect ER or golgi morphology (Figure S2A). Pseudo-cleavage furrows that formed properly and did not retract in Btsz-RNAi embryos reached their maximum length and were comparable in length to wild-type furrows (Figure S2B, C). However, the straightness of the furrows projecting into the cytoplasm were often abnormal in Btsz-RNAi embryos – furrows were more bent than in wild type, which could be observed across the embryo by comparing the outlines of pseudo-cleavage furrows at different depths (Figure S2D). Although F-actin levels were equivalent between control and Btsz-RNAi pseudo-cleavage furrow, the different morphology and stability and the absence of a change in ER/golgi morphology suggested that the mechanics or structure of these furrows is compromised.

### Btsz depletion leads to spindle collision and nuclear defects

Because pseudo-cleavage furrows are essential for preventing spindle collisions during each nuclear cycle, we assessed the integrity of the nuclear divisions in Btsz-depleted embryos. Using the microtubule plus-end-tracking protein CLIP170 fused with GFP to visualize spindles, we found that aberrant spindle morphology accompanied *Btsz* knock-down (Figure 3A, C). In Btsz-depleted embryos, spindles were often abnormally fused between neighboring nuclei (Figure 3A). This spindle collision phenotype has been noted in various conditions where proteins that regulate actin are disrupted (Dia, Arp2/3, Par1, APC2, Abl, Nuf, Rab11, etc.) (Grevengoed et al., 2003; Jiang and Harris, 2019; McCartney et al., 2001; Riggs et al., 2003; Rothwell et al., 1998; Webb et al., 2009; Zhang et al., 2018).

**Figure 3.**
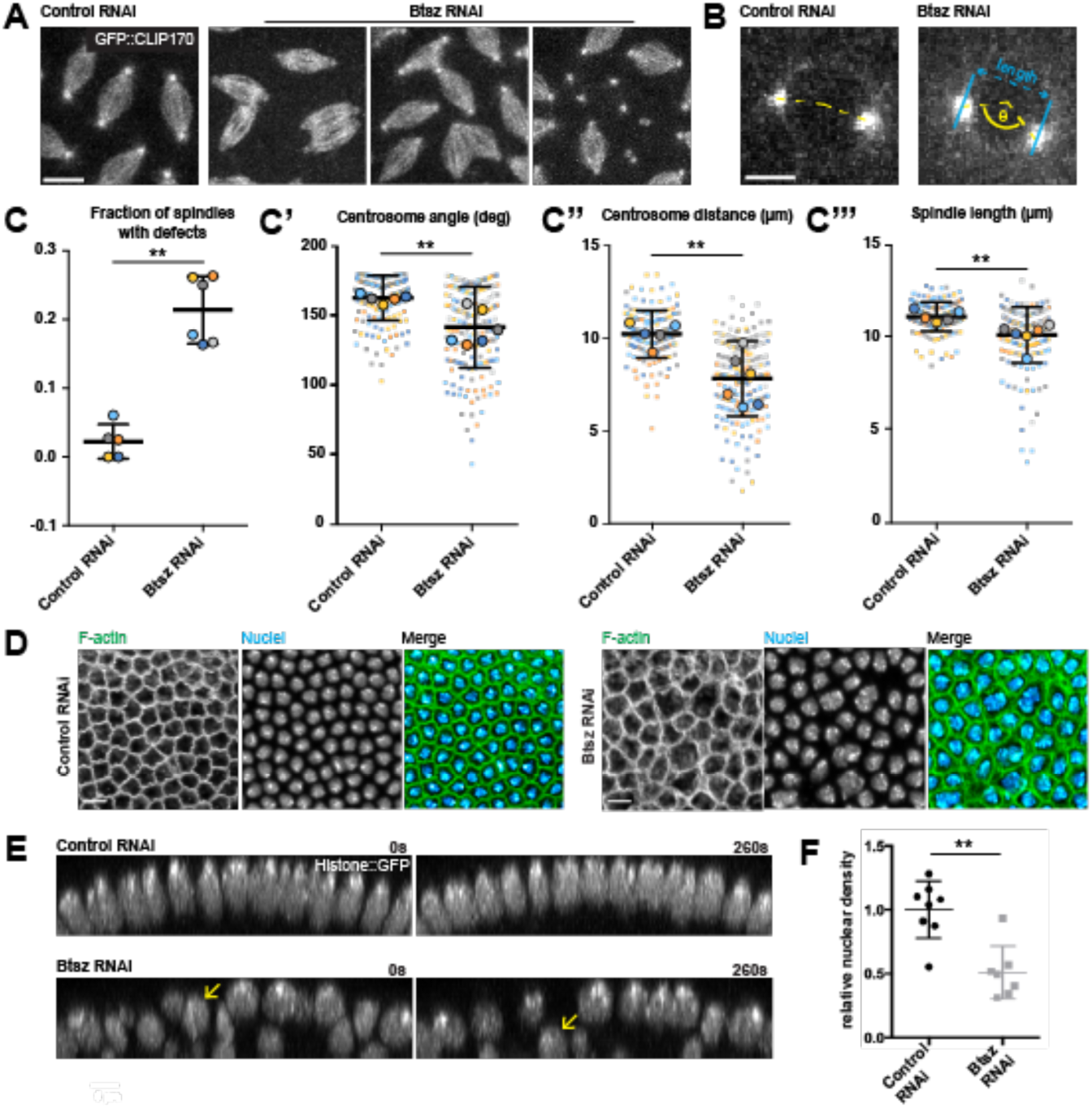
Btsz-depletion leads to spindle collision and nuclear defects. **(A, B)** Maximum projections of nuclear cycle 11 spindles and centrosomes visualized live using GFP::CLIP170 in control and Btsz-depleted embryos. Spindle collisions and aberrant centrosome morphology occur more frequently in Btsz RNAi embryos. Scale bars are 10 µm for spindles, 5 µm for centrosomes. **(C)** Fraction of spindles that have collided in control and Btsz RNAi embryos. Each large, bold circle is the fraction of occurrences in an embryo **(C’-C’’’)** Quantification of shortest distance between a centrosome pair (cyan line), angle of a vertex drawn between a centrosome pair (yellow dashed line), and spindle length. Each large, bold circle is the average of one embryo and smaller circles of corresponding color are the centrosome pairs or individual spindles used in the analysis. n = 5 embryos for control RNAi, n = 6 embryos for Btsz RNAi. p = 0.0043 for fraction with defects, 0.0087 for centrosome distance and angle, 0.0043 for spindle length, Mann-Whitney U test. **(D)** Single sagittal slice of control RNAi or Btsz RNAi embryos during early cellularization. F-actin was visualized using Phalloidin and nuclei was visualized using Hoechst in fixed samples. Both the actin cytoskeleton and nuclei are abnormal in Btsz RNAi embryos. **(E)** Orthogonal slice of nuclei visualized live using Histone::GFP at cellularization in control (top) or Btsz RNAi (bottom) embryos at 0s and 260s later. Yellow arrow indicates one nucleus at the surface of the Btsz RNAi embryo undergoing fallout. **(F)** Nuclear fallout occurs more frequently in Btsz-depleted embryos compared to the control. Each point is the number of nuclei over area (nuclear density) in one embryo. p = 0.0043, Mann-Whitney U test.

Proper actin cytoskeleton regulation is important for centrosome migration to opposite poles during nuclear division cycles (Cao et al., 2008). Centrosome separation was abnormal in Btsz-depleted embryos (Figure 3B) with centrosomes failing to migrate to 180° opposite each other prior to nuclear envelope breakdown more often than in wild-type embryos (Figure 3B, C’). Both the distance between centrosomes (Figure 3C’’) and spindle length (Figure 3C’’’) were shorter in Btsz-depleted embryos compared to wild-type embryos. These phenotypes were reminiscent of those observed in other actin and APC2 mutants (Cao et al., 2008; McCartney et al., 2001). Fixed imaging of both F-actin and nuclei during cellularization revealed heterogeneity in compartment sizes and nuclear morphology in Btsz-RNAi embryos, indicative of improper nuclear divisions that occurred earlier in the syncytium (Figure 3D).

Visualizing both F-actin and spindles confirmed that furrow breakages in the Btsz-RNAi background led to two sets of spindles in the same compartment (Figure S2E). These spindle abnormalities preceded massive nuclear fallout from the cell cortex during syncytial stages and preceding cellularization, observed through live imaging the nuclear marker Histone::GFP (Figure 3E, yellow arrows; Video 3). Quantification of relative nuclear density showed that there are fewer nuclei in Btsz-depleted embryos compared to the control (Figure 3F), likely a consequence of the pseudo-cleavage furrow defects (Figure 2C) and subsequent nuclear fallout. Overall, these results demonstrate a critical role for Btsz in embryo development, prior to its known role in epithelial integrity during gastrulation.

### Btsz has function(s) in the syncytium that do not depend on localizing Moesin

Prior work showed that Btsz functions with Moesin during gastrulation (Pilot et al., 2006b) and Moesin has been suggested to function in syncytial nuclear divisions (Vilmos et al., 2016). To determine how Btsz regulates syncytial divisions we examined Moesin and Btsz localization in the syncytial embryo. We first examined the localization of phospho-Moesin (pMoesin), the activated form of Moesin, using a previously published antibody (Carreno et al., 2008). pMoesin staining was not enriched in actin caps or pseudo-cleavage furrows (Figure 4A, Figure S2F). However the pMoesin antibody clearly labeled a perinuclear structure as has been described previously (Vilmos et al., 2009). We found this pMoesin-containing structure colocalized with Lamin B and was present at different cell cycle stages, in a manner that resembles Lamin B staining (Figure 4A). Because a nuclear excluded mutant of Moesin (fused to nuclear export sequence) also exhibits this staining pattern, we suggest that pMoesin, in part, is coating the nuclear envelope (Bajusz et al., 2021). BtszF is one of the shortest Btsz isoforms (Figure 1C), containing only the C2 domains that are necessary for proper localization and one additional exon. Because past work indicated that the C2 domains are responsible for Btsz localization (Pilot et al., 2006b), we visualized this tagged isoform to infer Btsz localization. Unlike pMoesin, BtszF localized to pseudo-cleavage furrows and plasma membrane infoldings from the actin caps (Tam and Harris, 2024) (Figure 4B). Thus, Btsz and pMoesin localize to different subcellular structures in syncytial embryos.

**Figure 4.**
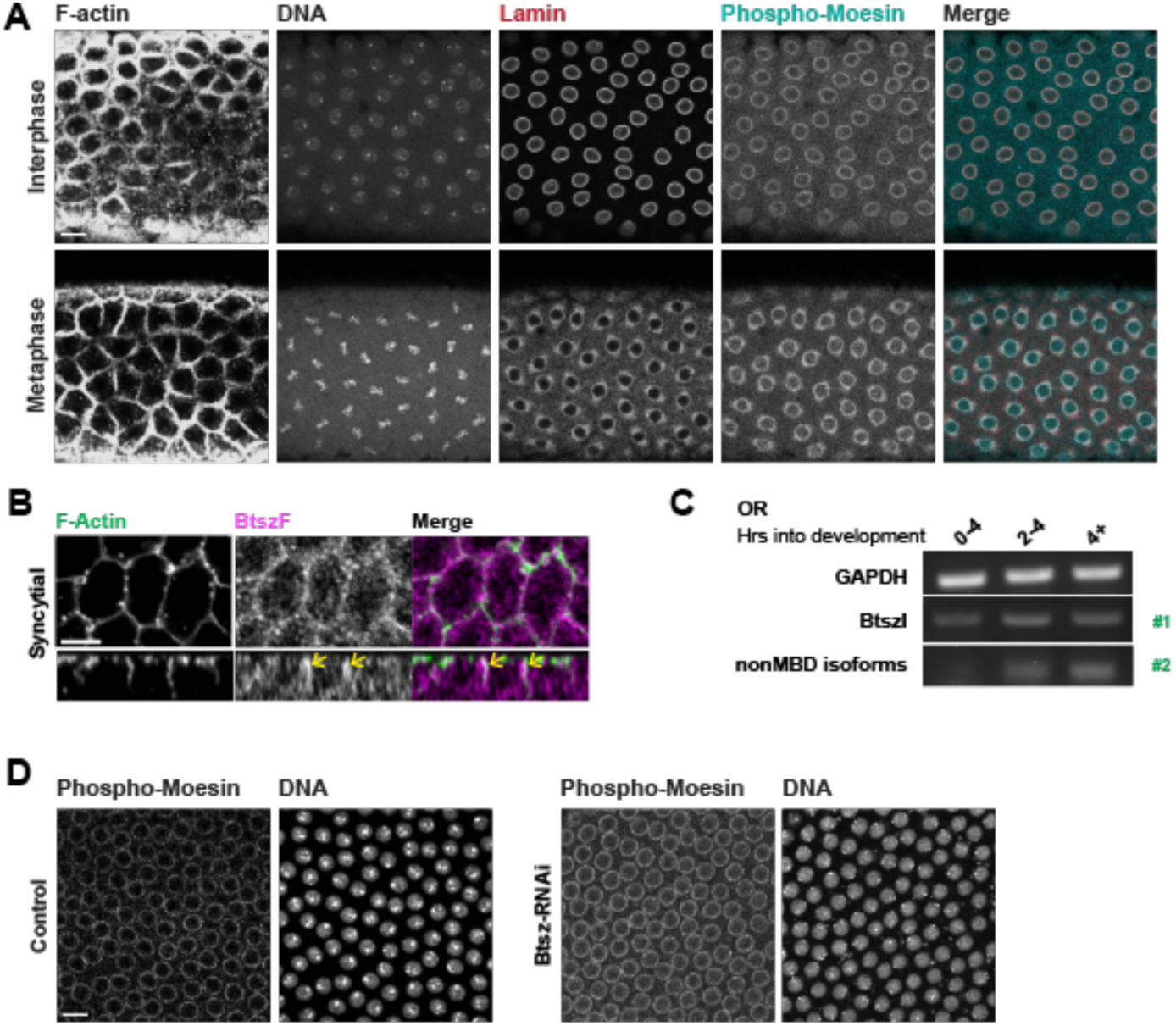
Btsz has function(s) in the syncytium that do not depend on localizing Moesin. **(A)** Optical section of the F-actin network, DNA, Lamin B, and phospho-Moesin localization in the wild-type syncytium at interphase and at metaphase. Phospho-Moesin localizes to perinuclear regions, closely overlapping with Lamin B. **(B)** Maximum projection of a top-down view of the F-actin network and Btsz isoform F localization in the syncytium. BtszF localizes to pseudo-cleavage furrows (yellow arrows). **(C)** Gel of RT-PCR products measuring isoform expression over developmental time. Both non-MBD isoforms and MBD isoforms are upregulated during the maternal to zygotic transition (MZT). Primer pair #1 detected BtszI and primer pair #2 detected non-MBD isoforms, as indicated in green. **(D)** Optical section of phospho-Moesin localization during interphase is similar for wild-type and Btsz-RNAi syncytium. Scale bars: A, D, 10 μM; B, 5 µm.

We next determined which Btsz isoforms were expressed in the early embryo. Consistent with what was previously reported, we found that Moesin Binding Domain (MBD, Exon8) isoforms and Exon 4-6-containing exons were present in the early embryo, consistent with prior work (Pilot et al., 2006b) (Figure S1A and B). Because BtszI has the MBD exon and most other exons, we distinguished between BtszI and non-MBD isoforms by using isoform specific primers (Figure 1C, green arrows). We found that non-MBD isoforms, as well as BtszI, increased in expression at the MZT ∼ 2-4 hr time point (Figure 4C). Detection of BtszI and non-MBD isoforms was inconsistent at 0-2 hours, suggesting that there is low or no expression of these isoforms at this stage. Overall, this data shows that MBD, non-MBD, and BtszI isoforms are expressed in the early embryo.

We next tested whether pMoesin localization is compromised in Btsz-RNAi embryos with syncytial defects. We found that Btsz-RNAi did not affect pMoesin intensity or localization, suggesting that the syncytial defects in Btsz-RNAi did not result from activated Moesin recruitment defects (Figure 4D). Taken together, the distinct localizations of Btsz and Moesin, the absence of Moesin at pseudo-cleavage furrows, failure of Btsz-RNAi to affect Moesin localization, and the presence of both MBD and non-MBD isoforms suggests that Bitesize has some functions that are Moesin independent.

### Btsz mutants suggest Moesin-independent function(s) during syncytial cell divisions

While we found that Bitesize expression was upregulated at cellularization (Figure 4C), Btsz-RNAi did not significantly affect cellularization furrow canal F-actin levels (Figure S3A and S3B). Cellularization was sometimes uneven in Btsz-RNAi embryos most often exhibiting a delay at the posterior pole (Figure S3C). Because perturbations that disrupt nuclear density result in significant asynchrony in cell cycle times (Deneke et al., 2019), it is likely that Btsz-RNAi cellularization phenotypes reflect earlier spindle collisions. However, abnormal nuclear density at cellularization (Figure 5C) gave us a robust metric to compare the severity of the nuclear fallout resulting from spindle collisions during syncytial divisions (Video 3).

**Figure 5.**
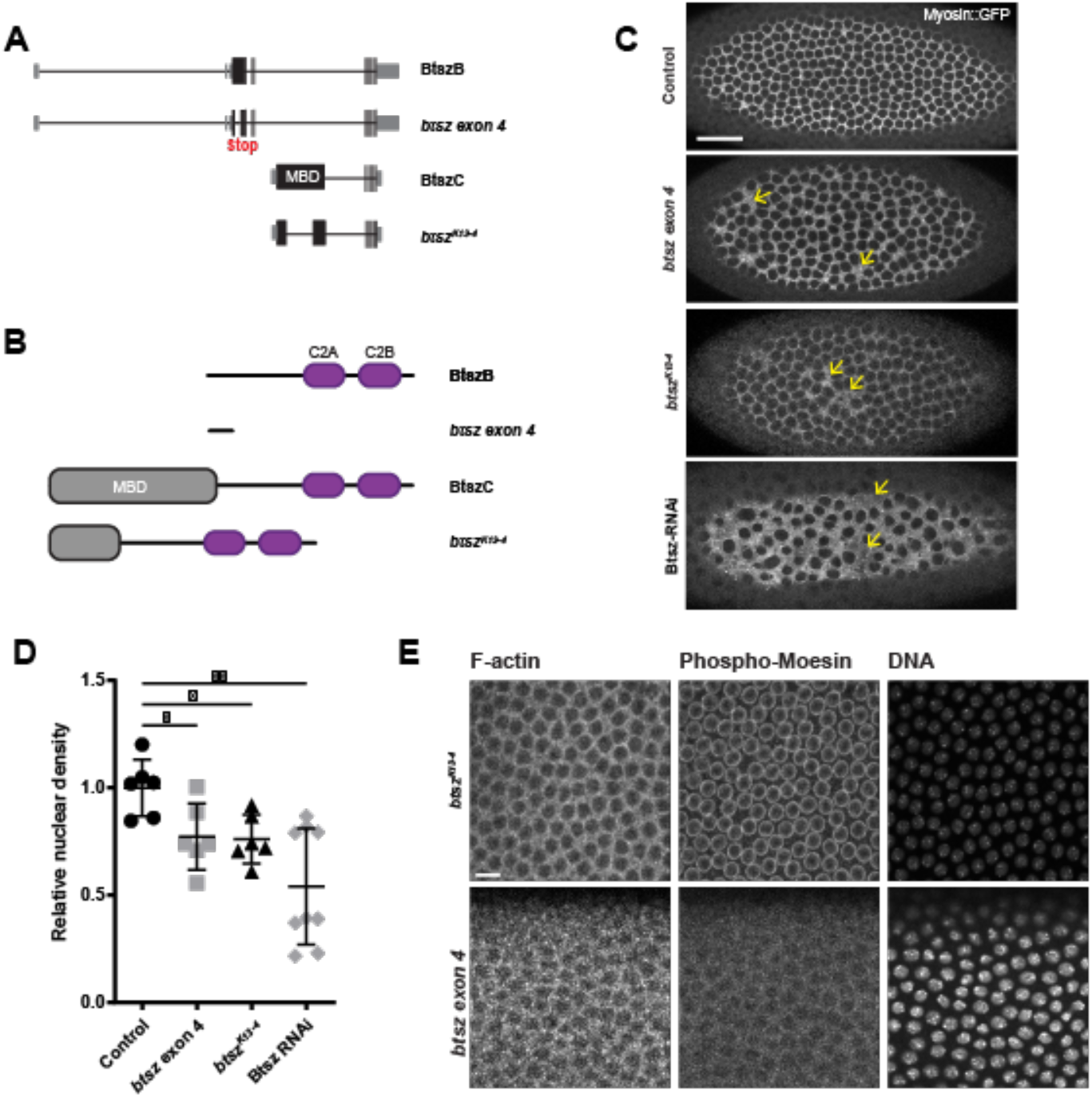
Non-MBD and MBD containing Btsz isoforms are important for development. **(A)** Schematic of Btsz isoforms B and C, along with Btsz mutants *btsz exon 4 (*which was generated in this study) and *btsz^K13-4^*. *btsz exon 4* has an early truncation in exon 4, and *btsz^K13-4^* has a deletion in exon 8. **(B)** Schematic of the protein products of the Btsz isoforms and mutants shown in (B). **(C)** Maximum projection images of cellularizing live embryos expressing Spaghetti squash::GFP (Sqh::GFP). Labeling myosin allows for visualization of the cytoskeletal network between nuclei enabling robust nuclear density determination. Yellow arrows indicate gaps where nuclei have dropped out that have been filled in by the myosin network. Scale bar is 25 µm. **(D)** Relative nuclear density in *btsz exon 4* and *btsz^K13-4^* mutants, compared to controls. p = 0.0260 for both comparisons, Mann-Whitney U test. **(E)** Grazing slices of cortical nuclei in syncytial embryo. F-actin was visualized using Phalloidin, nuclei was visualized using Hoechst, and Phospho-Moesin with anti-pMoe in fixed samples. Scale bar is 10 µm.

Because Btsz-RNAi embryos exhibited an obvious nuclear density phenotype and because live actin markers occasionally modified the furrow canal phenotypes (Spracklen et al., 2014), we visualized nuclear compartments using the non-muscle myosin II marker, Spaghetti squash::GFP (Sqh::GFP) (Figure 5C,D). Btsz-RNAi resulted in massive nuclear fallout, which was reflected in abnormal nuclear density by the time of cellularization, with the myosin network coating the membrane spanning the space between nuclei. Note that there were some more mildly affected embryos in the Btsz-RNAi data, which likely reflected incomplete knock-down.

To determine Btsz function in syncytial stages, we examined mutants that affect different sets of Btsz isoforms. The *btsz^K13-4^*mutant has a deletion in the MBD-containing exon 8, which affects BtszC and BtszD, as well as the longest isoform, BtszI (Serano and Rubin, 2003). Germline *btsz^K13-4^* mutant clones showed abnormal nuclear density (Figure 5D) and actin caps that appeared to have lost nuclei at syncytial stages (Figure S4), but did not affect pMoesin localization (Figure 5E). The *btsz^K13-4^*mutant affecting all MBD isoforms, including BtszI, but not recapitulating the pan-isoform RNAi knock-down, again argues for Moe-independent functions during syncytial development. If this were the case, we would expect other isoforms to also contribute to syncytial development.

Therefore, we used CRISPR/Cas9 to engineer a *btsz* mutant (*btsz exon 4*) with an early truncation present in the majority of non-MBD isoforms and BtszI (Figure 5A-B). The *btsz exon 4* mutant affected nuclear density and also did not fully recapitulate the pan-isoform knock-down (Figure 5C and 5D). Surprisingly, we found that the *btsz exon 4* mutant affects the nuclear localization of pMoesin. In summary, the full activity of Btsz requires both sets of MBD and non-MBD isoforms and the phenotype does not depend on an effect on pMoesin localization, suggesting that there are Moesin-independent functions for Btsz in the syncytium.

### Btsz C-terminus binds and bundles F-actin

Given the defects in actin-related processes exhibited by Btsz-RNAi and Btsz mutants, we hypothesized that Btsz, including non-MBD isoforms, may directly interact with F-actin. We examined Btsz function *in vitro* using the BtszB-C terminal fragment, which starts within exon 6 and contains the C2A and C2B domains shared by the majority of Btsz isoforms (Figure 6A). The BtszB C-terminal fragment does not contain the MBD. The purified C-terminal fragment of BtszB (BtszB-CT) demonstrated good expression and solubility.

**Figure 6.**
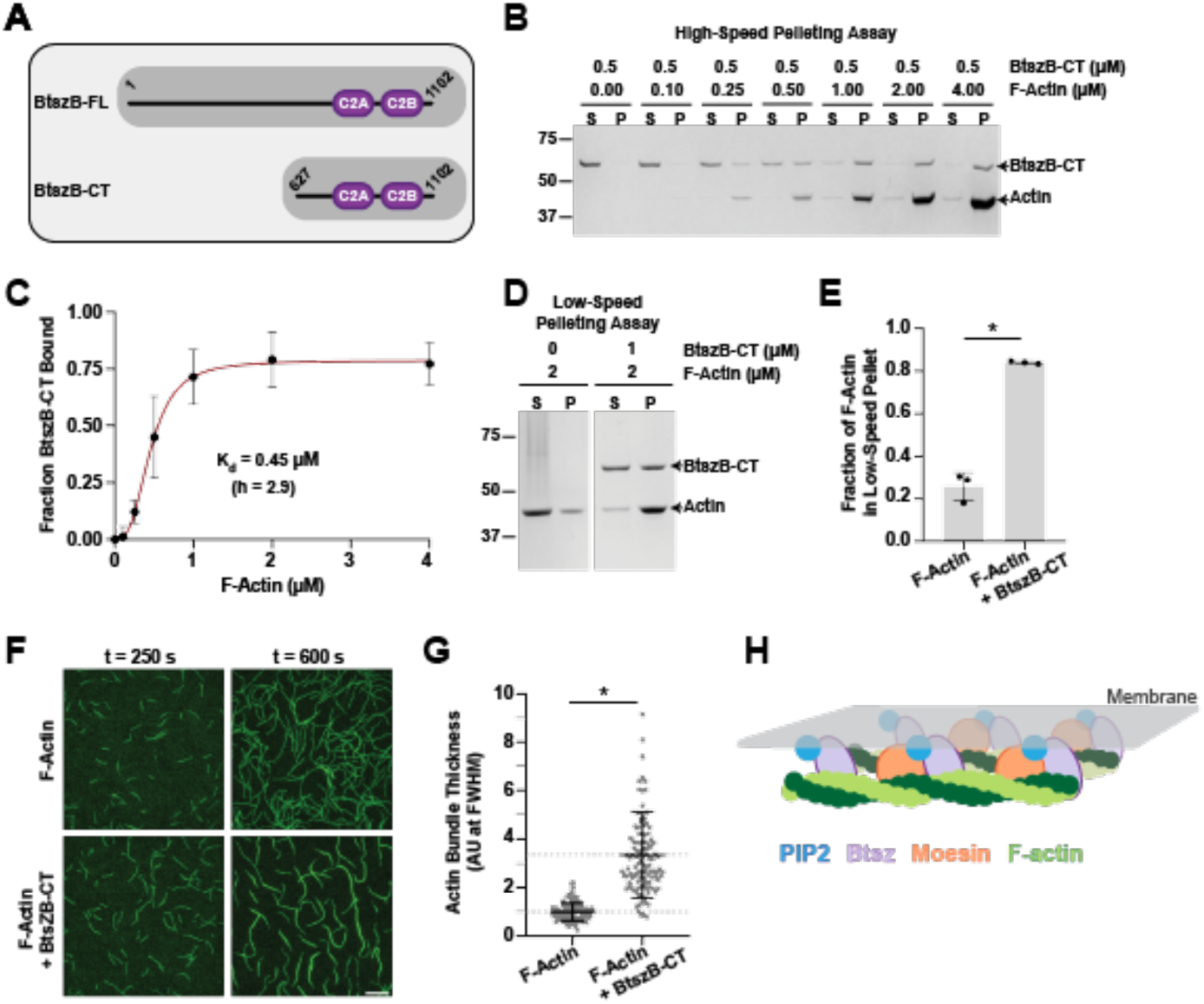
The C-terminus of BtszB cooperatively binds and bundles actin filaments. **(A)** Schematic of full length BtszB (BtszB-FL) for reference and the C-terminal fragment (BtszB-CT) that was used in the F-actin binding and bundling assays. **(B)** High-speed F-actin co-sedimentation assay varying the F-actin concentration (0.1 - 4 µM) in the presence of 0.5 µM BtszB-CT. F-actin and BtszB-CT samples were incubated together at room temperature for 30 min and then centrifuged (20 min at 25°C, 316k x g). **(C)** The fraction of BtszB-CT bound to actin filaments at increasing concentrations of F-actin was fit to a cooperative binding model with a Kd of 0.45 µM and a hill coefficient (h) of 2.9. Error bars are SD, n = 3. **(D)** Low-speed F-actin co-sedimentation assay using 2 µM F-actin with and without 1 µM BtszB-CT. F-actin and BtszB-CT were mixed together and incubated at room temperature for 30 min, then centrifuged (10 min at room temperature, 16k x g). **(E)** The fraction of F-actin pelleted with and without BtszB-CT. Error bars are SD, n = 3. p-value < 0.001, Student’s t-test. **(F)** Representative time-lapse images from TIRF microscopy reactions containing 2 µM G-actin (10% Alexa 488-labeled) polymerized into filaments 10-20 µm in length, and then buffer or 1 µM BtszB-CT was flowed into the reaction chamber at 300 s and monitored for an additional 300 s. Scale bar is 20 µm. **(G)** Filament/bundle thickness was assessed by measuring the fluorescence intensity at FWHM (full width at half maximum) from line segments drawn perpendicular to the filament or bundle. Fluorescence intensity values were normalized to the mean intensity value of the control reaction (2 µM F-actin). Horizontal dashed lines are at y = 1, mean 2 µM F-actin control reaction, and y = 3.4, mean 2 µM F-actin + 1 µM BtszB-CT. (nexperiments = 3, nline segment measurements for 2 µM F-actin = 41, 40, 42 and 1 µM BtszB-CT = 24, 35, 57. Error bars are SD, * is p-value < 0.001, Mann-Whitney U test). **(H)** Schematic of how Btsz non-MBD and MBD isoforms may interact with Moesin and/or F-actin at the membrane to bundle and organize the cytoskeleton.

We performed high-speed pelleting assays to test whether BtszB-CT binds to actin filaments and low-speed pelleting assays to test whether BtszB-CT bundles F-actin. In high-speed pelleting assays, a fixed concentration of BtszB-CT was mixed with variable concentrations of F-actin, and then the amount of BtszB-CT bound to F-actin was determined by analyzing the supernatant and pellet fractions on gels. With increasing concentrations of F-actin, BtszB-CT shifted from the supernatant to the pellet (Figure 6B). The data best fit a cooperative binding model with a binding affinity (K_d_) of 0.45 µM and a Hill coefficient (h) of 2.9 (Figure 6C). In low-speed pelleting assays, F-actin alone did not pellet significantly, but the addition of BtszB-CT shifted F-actin to the pellet (Figure 6D and E), suggesting that BtszB-CT organizes F-actin into higher order structures. These effects were confirmed independently through direct visualization in TIRF microscopy assays, where BtszB-CT organized F-actin into bundles with increased fluorescence and thickness (Figure 6F and G). Overall, these results show that BtszB-CT binds directly and cooperatively to F-actin (K_d_ = 0.45 µM) and organizes actin filaments into bundles, suggesting that all Btsz isoforms may bind and bundle F-actin.

## Discussion

Actin network organization must be tightly regulated for proper syncytial blastoderm development. Our data reveal a novel role for Synaptotagmin-like protein Btsz in organizing actin in the syncytial *Drosophila* embryo. We found that Btsz depletion resulted in defects reminiscent of the phenotypes caused by other mutants and perturbations that affect actin organization, including defective actin caps and pseudo-cleavage furrows (Afshar et al., 2000; Stevenson et al., 2002; Zallen et al., 2002). Although prior work implicated Moesin downstream of Btsz function, our evidence suggested that Btsz syncytial function was independent of Moesin: 1) pMoesin localization was peri-nuclear and not enriched at actin caps or pseudo-cleavage furrows, 2) Btsz-RNAi resulted in a dramatic pseudo-cleavage furrow and spindle collision phenotype, including abnormal, bent furrow structures, but did not affect pMoesin localization, 3) we showed that most isoforms are expressed in the early *Drosophila* embryo, including those lacking the MBD, 4) the *btsz^K13-4^* mutant germline clones, which affect all MBD-containing isoforms including BtszI, did not recapitulate the pan-isoform Btsz-RNAi depletion and did not affect pMoe localization, 5) the *btsz exon 4* mutant, which affected isoforms lacking the MBD and BtszI, had a nuclear density phenotype, and 6) we showed that a Btsz C-terminal fragment (lacking the MBD) cooperatively binds to F-actin and promotes F-actin bundling. This supports a model in which Btsz binds the membrane through its C2 domains and directly promotes actin bundling along the membrane. In the related Slac2 protein, Melanophilin, the actin binding domain (ABD) is also at the C-terminus, however, the Btsz C-terminus is more similar to Slp proteins (Kuroda et al., 2003). The identification of Btsz C-terminal actin binding and bundling activity shows a unique activity that extends to proteins of the Slp family. Future studies are needed to identify Btsz’ actin binding domain and to separate the many activities of Btsz, membrane binding, pMoesin recruitment, and actin binding/bundling.

### Btsz regulates actin organization at multiple stages of early *Drosophila* **embryo development**

Our *in vitro* data indicate that the Btsz C-terminus binds and bundles F-actin, shedding light on how Btsz may regulate actin *in vivo* in a complementary manner to previously published role through Moesin (JayaNandanan et al., 2014; Pilot et al., 2006b; Zhang et al., 2018). Our data suggests that many isoforms could bind and bundle F-actin and moreover we did not see Moesin localization perturbed in all conditions that caused defects. In the syncytium, Btsz could promote actin filament bundles generated by the formin Diaphanous (Jiang and Harris, 2019). BtszF contains the C2 domains and localizes to syncytial pseudo-cleavage furrows; because all Btsz isoforms contain the C2 domains responsible for membrane localization, it is possible that all isoforms localize to pseudo-cleavage furrows. Btsz bundling may stabilize actin at the pseudo-cleavage furrows and prevent their recession, as furrows in Btsz-depleted embryos often fail and disappear. We also observe that the psuedo-cleavage furrows of Btsz-depleted embryos appear to become more undulatory over time compared to controls, although higher resolution imaging is required to determine whether this is the result of a more global perturbation in F-actin organization in these embryos (Figure 2C, Figure S2C). The development of undulations may be indicative of a mechanical instability (in addition to the temporal instability discussed above), which could arise due to a lack of actin organization. During oogenesis, the *Drosophila* Fascin homolog, Singed, is required for actin bundling when rapid actin polymerization takes place to provide structural integrity for these bundles (Cant et al., 1994). Btsz may function in a similar way, stabilizing F-actin bundles against disassembly during a time of rapid actin remodeling during syncytial nuclear divisions. However, an important next step will be to make mutants that separate Btsz actin binding and bundling activities and determine their in vivo function.

Btsz isoforms are upregulated at the MZT. Zygotic gene activity is present as early as cycle 10, where transcription is required for patterns of nuclear density (Blankenship and Wieschaus, 2001). The expression of many actin remodeling proteins are developmentally upregulated at the MZT, which is necessary for cellularization.

These include Bottleneck, Nullo, Slam, Dunk, and Serendipity-α (Sry-α), are developmentally upregulated at the MZT (He et al., 2016; Krueger et al., 2019; Lecuit et al., 2002; Postner and Wieschaus, 1994; Schejter and Wieschaus, 1993; Simpson and Wieschaus, 1990; Sokac and Wieschaus, 2008; Wenzl et al., 2010; Zheng et al., 2013). Unlike Btsz, these genes have a simple structure and lack complex splicing patterns.

Unlike *nullo* or *sry-α*, Btsz does not affect F-actin levels in furrow canals during cellularization (Figure S3A, B) (Postner and Wieschaus, 1994; Simpson and Wieschaus, 1990), suggesting that Btsz functions to prevent spindle collisions and nuclear fallout during syncytial divisions.

Actin bundles are also a key component of adherens junctions (AJs), which are critical for cell-cell adhesion. The dynamic coupling of the F-actin network to adherens junctions is necessary for propagating forces across cells (Gorfinkiel and Arias, 2007; Jodoin, 2015; Maître et al., 2012; Martin et al., 2010; Roh-Johnson et al., 2012). E-cadherin (E-cad) is an important component of AJs, and its localization is stabilized by F-actin. Non-junctional E-cadherin localizes to pseudo-cleavage furrows during metaphase so it is possible that Btsz also interacts with E-cadherin in the syncytium (Rikhy et al., 2015). When Btsz is disrupted, so is actin organization which in turn destabilizes E-cadherin, leading to defects in tissue integrity (Pilot et al., 2006b). It is possible that multiple Btsz isoforms with distinct activities could collaborate bind and bundle F-actin to organize actin at these junctions. Overall, we speculate that Btsz is a multifunctional cytoskeletal organizer, coupling membrane-cortex anchoring and F-actin bundling activities.

### Possible functions for Btsz in membrane trafficking

An important question that remains unanswered is whether Btsz plays a role in *Drosophila* regulating membrane trafficking like mammalian Slps. Our work here focused primarily on Btsz’s role in actin organization, yet actin regulation and membrane regulation may be interrelated. For instance, the Nuclear fallout (Nuf) protein is required for actin and membrane remodeling during the syncytial divisions through its role in regulating trafficking (Riggs et al., 2003). In *nuf* mutants, pseudo-cleavage furrows are defective leading to spindle collisions and the ‘nuclear fallout’ phenotype (Rothwell et al., 1998).

One surprising result in our work was that *btsz exon 4* mutant disrupted pMoesin’s peri-nuclear localization, while the pan-isoform Btsz-RNAi knockdown did not. Considering the close coupling between actin and membrane remodeling, it is possible that the nuclear fallout phenotype is, in part, due to defects in vesicle trafficking. Moreover, BtszF is present in puncta (Figure 4B) which may indicate a potential role in membrane trafficking and/or Btsz localization at plasma membrane invaginations associated with exocytosis (Tam and Harris, 2024). Interestingly, the N-terminus of several Btsz isoforms contains a putative zinc finger domain similar to those that interact with the Rab GTPases. The *btsz exon 4* mutant truncates protein products after this zinc finger domain, which could have an effect on vesicle trafficking, pMoesin’s recruitment to the peri-nuclear structure, or Moesin phosphorylation. Future work is required to dissect the nature of pMoe nuclear localization and its regulation by Btsz. However, our work has shown a Moesin-independent function for some Btsz isoforms .

## Materials and Methods

### Fly stocks and genetics

Fly stocks and crosses used in this study are listed in Supplementary File 1. For crosses involving maternal gene depletion and overexpression, flies expressing maternal Gal4 drivers were crossed to UAS-driven constructs and maintained at 25°C. Nonbalancer F1 females were crossed to nonbalancer F1 males, and F2 embryos were used for imaging.

For the *btsz exon 4* mutant, a deletion was generated at the endogenous Btsz locus using CRISPR-Cas9 as previously described (Gratz et al., 2015). Two 15 base pair (bp) gRNAs targeting sites in Btsz exon 4, roughly 500 bps apart, were identified using flyrnai.org/crispr and cloned into the pCFD5 plasmid. This vector was constructed and packaged by VectorBuilder and the vector ID is VB200707-1018qzm, which can be used to retrieve further information about the vector on vectorbuilder.com. The plasmid was injected into nanos-Cas9-expressing embryos by BestGene Inc (Chino Hills, California). Surviving adults were crossed to Dr/TM3 flies and the deletion was screened for using PCR. Flies with successfully a generated deletion were sequenced for further analysis. For crosses involving *btsz exon 4*, crosses were set up using two *btsz exon 4* lines of independent CRISPR alleles.

For crosses involving *btsz^K13-4^*, the FLP-DFS technique was used to generate mutant germline clones (Chou and Perrimon, 1992). We used *btszK13-4* alone and sqh::GFP; *btsz^K13-4^* mutant stocks. Briefly, hsFLP;; Dr/TM3, Sb females were crossed to FRT82B ovoD1/TM3, Sb males. hsFLP/Y;; FRT 82B ovoD1/Dr males were crossed to *btsz^K13-4^* FRT/TM3, Sb or sqh::GFP; *btsz^K13-4^* FRT/TM3, Sb females. Larvae from this cross were heat shocked for 2 hrs at 37°C for 3-4 days. hsFLP; + or sqh::GFP*; btsz^K13-4^* FRT/*ovoD* FRT females were crossed to OregonR males to collect embryos resulting from the *btsz^K13-4^*germline clones.

The pUASt-mCherry::BtszB fly line was constructed using the coding sequence of BtszB (generous gift from Thomas Lecuit, Collège de France). The vector used to overexpress mCherry::Btsz in our study was constructed and packaged by VectorBuilder (Chicago, Illinois). The vector ID is VB211103-1199vfw. This vector was injected into attP40 flies and surviving adults were crossed to Sp/CyO flies. Presence of the vector was screened for by presence of red eye color and confirmed with subsequent PCR and sequencing.

### Live and fixed imaging

Crosses were maintained at 25°C unless otherwise stated. Embryos were collected from apple-juice agar plates and embryos were staged using Halocarbon 27 oil. Embryos were then dechorionated in 50% bleach (Chlorox) for 2 minutes and rinsed twice with water.

For live imaging, embryos were mounted on a slide with embryo glue (Scotch tape dissolved in heptane). No. 1.5 coverslips were placed on the slide on either side of the mounted embryos and a No.1 coverslip was used to cover the embryos and create a chamber. This chamber was filled using Halocarbon 27 oil.

For fixed imaging involving Phalloidin staining, embryos were fixed using 4% paraformaldehyde (Electron Microscopy Sciences) in 0.1M phosphate buffer at pH 7.4 with 50% heptane (Alfa Aesar) and manually devitellinated. Embryos were washed in 0.01% Triton X-100 in PBS (PBS-T), incubated in primary antibodies followed by secondary antibodies (Supplementary File 1), and mounted onto a glass slide using AquaPolymount (Polysciences).

Live and fixed images were taken using a Zeiss LSM 710 point scanning confocal microscope with a 40x/1.2 C-Apochromat water objective lens or 63x/1.4 Plan-Apochromat oil objective lens, an argon-ion, 405 nm diode, 594 nm HeNe, 633 HeNe laser, and Zen software. Pinhole settings ranged from 1 to 2.5 Airy units. For two-color live imaging, band-pass filters were set at ∼410-513 nm for Hoechst, ∼490–565 nm for GFP, and ∼590–690 nm for mCherry (mCh). For three-color imaging, band-pass filters were set at ∼480–560 nm for Alexa Fluor 488, ∼580–635 nm for Alexa Fluor 568, and ∼660–750 nm for Alexa Fluor 647.

### Image processing and analysis

Figure images were processed in FIJI or Matlab R2019b. For actin cap area and nuclear fallout quantifications, images were run through a segmentation analysis (detailed below) in Matlab. For all figures, brightness and contrast were adjusted linearly. For Figures 2, 3, and 6, images were processed in FIJI with a Gaussian blur of 0.5. For Figures 7 and 4B, images were processed in FIJI with a Gaussian blur of 0.5 and using Subtract Background with a rolling ball radius of 50.0 pixels.

### Actin cap and cellularization furrow quantifications

#### Actin cap area and nuclear fallout

Embryos expressing mCherry::Moe were used to visualize actin caps. To stage-match embryos, we scored the size and number of actin caps. We measured actin cap area during NC12 past the expansion phase of cap growth to minimize gaps in the actin network. This was roughly 4 minutes before pseudo-cleavage furrows reach their maximum depth during NC12. A maximum intensity projection (MIP) at this time point was then used in a segmentation pipeline in MATLAB detailed below and manual corrections were made in FIJI. To count nuclei, MIPs of embryos expressing Sqh::GFP were segmented using the same Matlab procedure and FIJI was used to measure embryo area in the given field of view and to count nuclei.

1. The intensity of each pixel was scaled up by a factor of 1.25.
2. Ridge-like structures in the image were enhanced with FrangiFilter2D using the default parameters (Dirk-Jan Kroon (2022). Hessian based Frangi Vesselness filter (https://www.mathworks.com/matlabcentral/fileexchange/24409-hessian-based-frangi-vesselness-filter), MATLAB Central File Exchange); based on (Frangi et al., 1998).
3. The resulting image was blurred via a Gaussian blur of sigma 4-10 depending on the image using Matlab’s imgaussfilt() function.
4. Seeds for a watershed segmentation were automatically determined using the imregionalmin() function on the blurred, background-subtracted image.
5. The ridge-enhanced image from step 2 was blurred with a median filter of size 11 pixels, then the seeds were imposed as local minima on this image.
6. The watershed() function was used on this image to generate lines corresponding to the boundaries between actin caps.

#### Furrow actin intensity

Cellularizing embryos were fixed and imaged to calculate the ratio of actin intensity in the apical to basal regions in the furrows. FIJI was used to measure intensity. A ratio of intensity was taken of the most apical and basal 2 µm of the furrows. Intensities were normalized by rescaling values between 0 and 1 using (I-I_min_)/(I_max_-I_min_).

### Molecular biology

For RNA extraction, thirty embryos were collected for each stage analyzed and ground up in TRIzol reagent (Life Technologies, Inc.) using a motorized pestle mixer. mRNA was extracted using TRIzol and chloroform. The aqueous phase was transferred into a fresh tube and extraction continued using the Monarch RNA Cleanup Kit (NEB). The extracted mRNA was treated with TURBO DNA-*free*^TM^ (Invitrogen) and cDNA was produced using Invitrogen SuperScript™ III First-Strand Synthesis System. Subsequent PCR reactions were run with primers listed in (Supplementary File 1).

### Purification of BtszB

Vectors of full-length BtszB and BtszB fragments were generated for protein expression and purification (ABclonal Science Inc.). Full-length Btsz isoform B was insoluble in bacteria and so we used the baculovirus system to express Btsz-B in Sf9 cells and purified protein from those (ABclonal Science Inc).

### Purification and Labeling of Actin

Rabbit skeletal muscle actin was purified from acetone powder (Spudich and Watt, 1971) generated from frozen ground hind leg muscle tissue of young rabbits (PelFreez). Lyophilized acetone powder stored at -80°C was mechanically sheared in a coffee grinder, resuspended in G-buffer (5 mM Tris-HCl [pH 7.5], 0.5 mM Dithiothreitol [DTT], 0.2 mM ATP, 0.1 mM CaCl_2_), and then cleared by centrifugation for 20 min at 50,000 × g. The supernatant was filtered through Grade 1 Whatman paper, then the actin was polymerized by the addition of 2 mM MgCl_2_ and 50 mM NaCl and incubated overnight at 4°C. The final NaCl concentration was adjusted to 0.6 M to strip residual actin-binding proteins, incubated at 4°C for 1 h, and then the F-actin was pelleted by centrifugation for 150 min at 361,000 × g. The pellet was solubilized by dounce homogenization and dialyzed against 1 L of G-buffer at 4°C (three consecutive times at 12–18 h intervals). Monomeric actin was then precleared at 435,000 × g and loaded onto a S200 (16/60) gel-filtration column (GE Healthcare) equilibrated in G-Buffer. Peak fractions containing actin were stored at 4°C.

To fluorescently label actin, G-actin was polymerized by dialyzing overnight against modified F-buffer (20 mM PIPES [pH 6.9], 0.2 mM CaCl_2_, 0.2mM ATP, 100 mM KCl) (Shekhar, 2017). F-actin was incubated for 2 h at room temperature with Alexa 488 NHS ester dye (Life Technologies) at a final molar concentration 5 times in excess of actin concentration. F-actin was then pelleted by centrifugation at 450,000 × g for 40 min at 4°C. The pellet was homogenized with a dounce and dialyzed overnight at 4°C against 1 L of G-buffer (three consecutive times at 12–18 h intervals). Next, the solution was centrifuged at 450,000 × g for 40 min at 4°C. The supernatant was collected and loaded onto a S200 (16/60) gel-filtration column (GE Healthcare) equilibrated in G-Buffer. Peak fractions containing labeled G-actin were pooled and the concentration and labeling efficiency was determined by measuring the absorbance at 290 nm and 494 nm. Molar extinction coefficients used were as follows: ε_290_ actin = 26,600 M^-1^ cm^-1^ and ε_494_ Alexa 488 = 71,000 M^-1^ cm^-1^. The labeled G-actin was stored in G-Buffer + 50% glycerol at -20°C and then dialyzed back into G-Buffer (no glycerol) for experiments.

### High-Speed and Low-Speed Pelleting (Co-Sedimentation) Assays

F-actin was prepared by polymerizing 20 µM of G-actin with initiation mix (final concentration: 2 mM MgCl_2_, 0.5 mM ATP, 50 mM KCl) overnight at room temperature. For high-speed pelleting assays, different concentrations of F-actin (0, 0.1, 0.25, 0.5, 1, 2, 4 µM) were incubated with 0.5 µM BtszB-CT for 30 min at room temperature.

Reactions were then centrifuged at 316,000 × g for 20 min at 25°C using a tabletop Optima TLX Ultracentrifuge (Beckman Coulter). Supernatant and pellet fractions were analyzed on Coomassie stained gels, and quantified by scanning densitometry using a ChemiDoc Imaging System (BioRad). In calculating the fraction of BtszB-CT bound to F-actin in each reaction, the BtszB-CT that pelleted nonspecifically (without F-actin) was subtracted. Fraction of BtszB-CT bound at different concentrations of F-actin was plotted, and fit to a Hill Equation using Prism (GraphPad). For low-speed pelleting assays, 2 µM of F-actin was incubated with or without 1 µM BtszB-CT for 30 min at room temperature. Reactions were then centrifuged at 16,000 × g for 10 min at room temperature using a tabletop centrifuge (Eppendorf). Supernatant and pellet fractions were analyzed on Coomassie stained gels and quantified as above. Fraction of F-actin pelleted +/-BtszB-CT was plotted and compared statistically using a Mann-Whitney U test from Prism (GraphPad).

### F-Actin Bundling TIRF Microscopy Assays and Image Analysis

Glass coverslips (60 × 24 mm; Thermo Fisher Scientific) were first cleaned by sonication in 1% Hellmanex™ III (Sigma-Aldrich) glass cleaner for 60 min at room temperature, followed by successive sonications in 100% ethanol for 60 min and 1 M KOH for 20 min. Coverslips were then washed extensively with H_2_O and dried in an N_2_ stream. The cleaned coverslips were coated with 2 mg/ml methoxy-polyethylene glycol (PEG)-silane MW 2000 and 2 µg/ml biotin-PEG-silane MW 3400 (Laysan Bio, Arab, AL) in 80% ethanol, pH 2.0, and incubated overnight at 70°C. Flow cells were assembled by rinsing PEG-coated coverslips with water, drying with N_2_, and adhering to µ-Slide VI0.1 (0.1 × 17 × 1 mm) flow chambers (Ibidi, Fitchburg, WI) with double-sided tape (2.5 cm × 2 mm × 120 µm) and 5-min epoxy resin (Devcon, Danvers, MA). Before each reaction, the flow cell was incubated for 1 min with 1% bovine serum albumin (BSA) in HEKG5 buffer (20 mM HEPES, pH 7.4, 1 mM EDTA, 50 mM KCl, and 5% glycerol), and then equilibrated with TIRF buffer (10 mM imidazole, pH 7.4, 50 mM KCl, 1 mM MgCl_2_, 1 mM ethylene glycol-bis(β-aminoethyl ether)-N,N,N′,N′-tetraacetic acid (EGTA), 0.2 mM ATP, 10 mM DTT, 15 mM glucose, 20 µg/ml catalase, 100 µg/ml glucose oxidase) plus 0.5% methylcellulose (4000 cP). Finally, 2 µM G-actin (10% Alexa 488–labeled on Cys 374) polymerized to 10 - 20 µm, washed with TIRF buffer and then 1 µM BtszB-CT was flowed into the reaction chamber 300 s after initiation of actin assembly. Upon flowing wash TIRF buffer and BtszB-CT into the TIRF chamber, filament often would move around but did not get flushed out because of the crowding agent (methylcellulose) in the buffer.

Single-wavelength time-lapse TIRF imaging was performed on a Nikon-Ti2000 inverted microscope equipped with a 150-mW argon laser (Melles Griot), a 60 × TIRF objective with a numerical aperture of 1.49 (Nikon Instruments), and an electron-multiplying charge coupled device (EMCCD) camera (Andor Ixon, Belfast, Ireland). One pixel was equivalent to 143 × 143 nm. Focus was maintained by the Perfect Focus system (Nikon Instruments). TIRF microscopy images were acquired every 5 s and exposed for 100 ms for at least 10 min using imaging software Elements (Nikon Instruments, New York, NY).

Images were analyzed in FIJI version 2.0.0-rc-68/1.52e (National Institutes of Health, Bethesda, MD). Background subtraction was conducted using the rolling ball background subtraction algorithm (ball radius, 5 pixels). To measure actin filament bundling, the line segment tool was used to draw a line perpendicular to actin filaments/bundles in the FOV. The intensity profile of the line segment was fit to a 2-dimensional Gaussian curve in FIJI. The intensity at full-width half-max (FWHM) for each line trace was recorded at 300 s into the reactions (where BtszB-CT was flowed in at time 0). All individual intensity measurements were normalized to the average actin filament/bundle intensity from the control reaction (2 µM F-actin). A Mann-Whitney U statistical test was performed to determine bundling activity between the control reaction (2 µM F-actin) and the 1 µM BtsB-CT reaction.

## Statistical analysis

All statistical analyses were carried out using GraphPad Prism or MATLAB.

## Acknowledgments

We thank members of the Martin lab for helpful discussions and comments on the manuscript. We thank Jonathan Jackson for writing the segmentation pipeline code. We thank J. Solitro, Z. Li, and J. Eskin for their assistance with experiments. We are grateful to Goode lab members for assistance with actin purification and labeling. Finally, we thank the Bloomington Stock Center and the TRiP at Harvard Medical School (National Institutes of Health/National Institute of General Medical Sciences R01-GM084947) for providing fly stocks used in this study. This research was supported by a grant from the National Institutes of Health to B.L. Goode (R35 GM134895). This work is also supported by National Institutes of Health grants R01 GM105984 and R35 GM144115 to A.C. Martin.

## Figures and Figure Legends

**Supplemental Figure 1.**
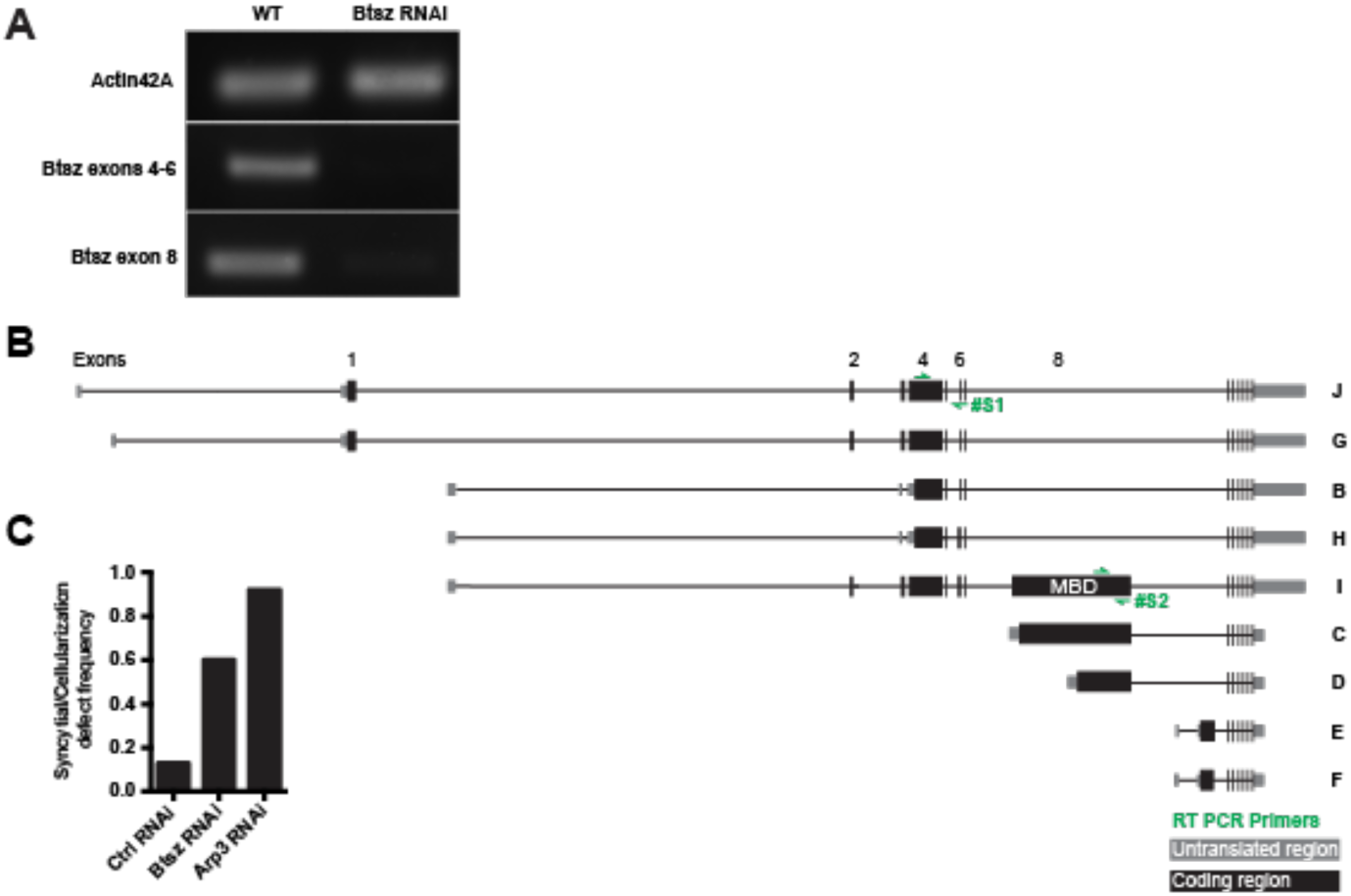
**(A)** RT-PCR Gel of Btsz cDNA showing maternal knockdown of Btsz in the early embryo. Primers used are shown in S1B. Primer pair S1 was used for Btsz exons 4-6 and pair S2 was used for Btsz exon 8, the MBD. **(B)** Schematic of the nine splice isoforms of the Btsz protein. Letters on the right-hand side denote the isoform name. Location of RT-PCR primer pairs #S1 and #S2 for exons 4-6 and exon 8, respectively, are shown with green arrows. **(C)** Frequency of control RNAi, Btsz RNAi, and Arp3 RNAi embryos with defects during the syncytial or cellularization stages. 4 out of 30 control, 25 out of 42 Btsz RNAi, and 12 out of 13 Arp3 RNAi embryos displayed defects.

**Supplemental Figure 2.**
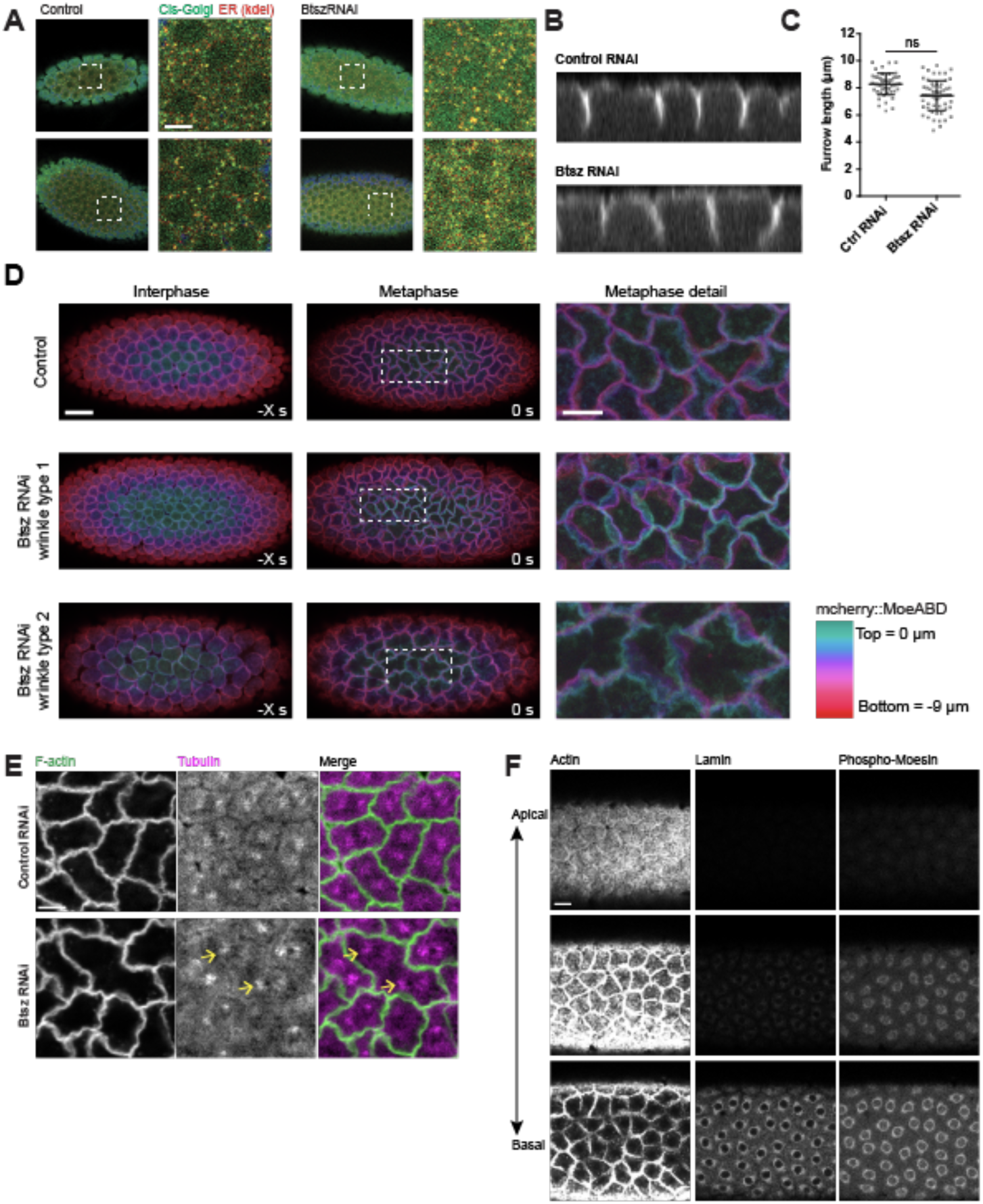
**(A)** Maximum intensity projections of cortical grazing slices of control and Btsz-RNAi embryos stained for ER (anti-KDEL) and Cis-golgi (anti-gm130). Morphology and overlap between stained compartments were not disrupted in Btsz-RNAi. Scale bar = 5 µm. **(B)** Cross section of pseudo-cleavage furrows during syncytial blastoderm development. Actin was visualized live with the mCherry::MoesinABD marker. **(C)** Pseudo-cleavage furrows that form normally in Btsz RNAi embryos are not significantly different in length compared to the wildtype. Each point is one furrow from n = 6 embryos for both control RNAi and Btsz RNAi. **(D)** Intensity projection of mCherry::MoesinABD with different depths encoded by color (right, colormap). Control embryos have tight distribution of colors, while Btsz-RNAi embryos often have dispersed color signals indicating tilted or wavy pseudo-cleavage furrow morphology. **(E)** Slice of fixed wildtype and Btsz RNAi embryos during metaphase. F-actin was visualized using Phalloidin and tubulin was visualized using an α-tubulin antibody. Pseudo-cleavage furrows that have receded lead to two sets of spindles in one compartment (yellow arrows). Scale bar is 5 µm. **(F)** Grazing sections of the F-actin network (Phalloidin), Lamin B, and phospho-Moesin in the wild-type syncytium at metaphase at dfferent depths. Phospho-Moesin localizes to perinuclear regions, closely overlapping with Lamin B and is absent from F-actin caps and pseudo-cleavage furrows.

**Supplemental Figure 3.**
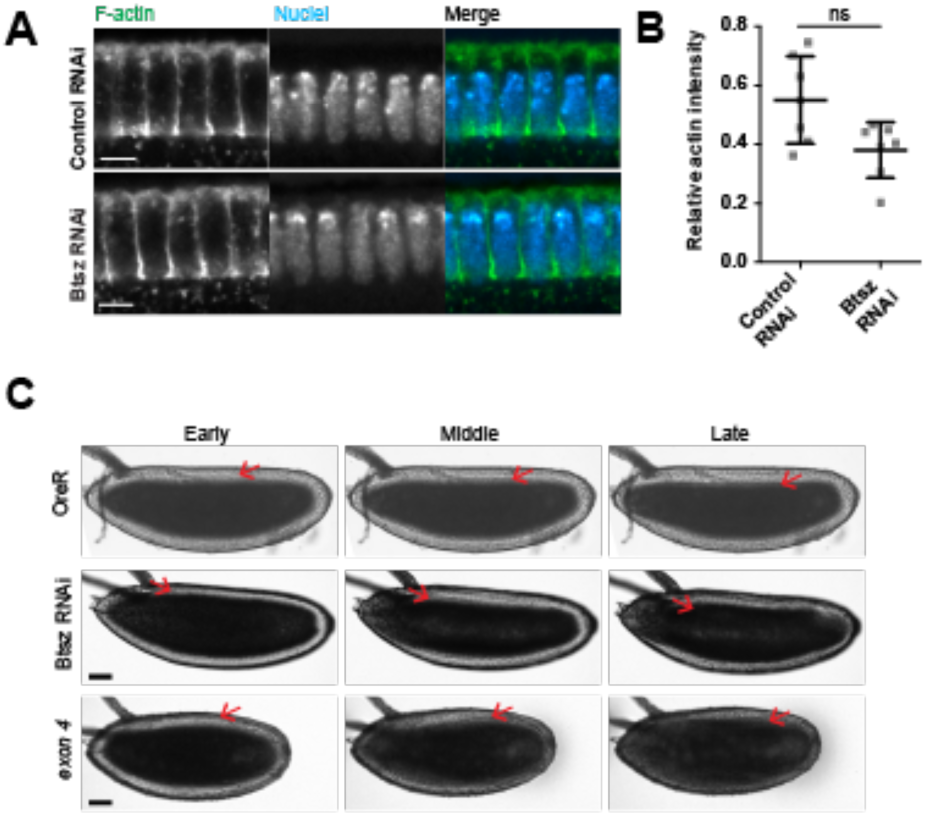
**(A)** Cross section of fixed control RNAi or Btsz-RNAi embryos during cellularization. F-actin was visualized using Phalloidin and nuclei were visualized using Hoechst. Scale bar = 5µm. **(B)** There is no significant difference between the relative F-actin intensity (ratio of actin intensity in the apical region to that of the furrow canals) in the wildtype and Btsz-RNAi embryos. Mann-Whitney U test. **(C)** Brightfield images of control, btsz-RNAi, and btsz[exon4] mutant embryos undergoing cellularization. Red arrows indicate cellularization front. Note that cellularization front is often uneven in Btsz-RNAi and btsz mutant embryos, likely due to nuclear fallout and density defects.

**Supplemental Figure 4.**
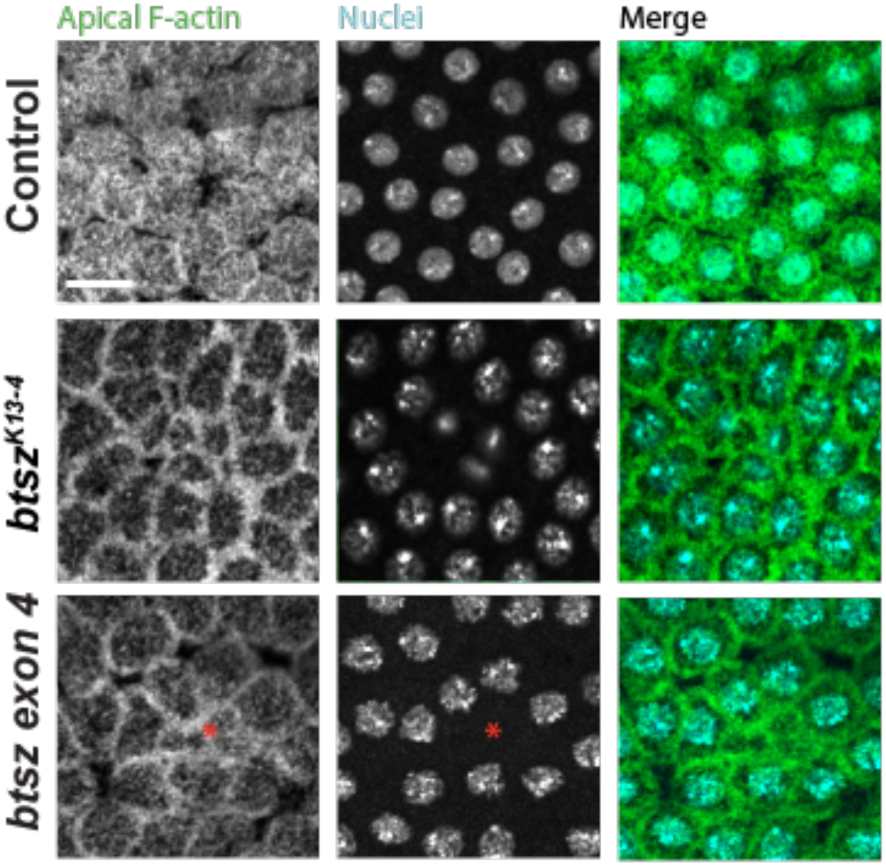
Btsz mutants have syncytial embryos defects. Images are grazing sections showing cortical F-actin (phalloidin staining) and nuclei (Hoechst staining) in control and btsz[k13-4] and btsz[exon4] mutants. Note that mutants exhibit regions that lack nuclei under actin caps (red asterisk), similar to the nuclear fallout phenotype of Btsz-RNAi. Scale bar = 10 µm.

